# Mosaic antimicrobial resistance/virulence plasmid in hypervirulent ST2096 *Klebsiella pneumoniae* in India: The rise of a new superbug?

**DOI:** 10.1101/2020.12.11.422261

**Authors:** Chaitra Shankar, Karthick Vasudevan, Jobin John Jacob, Stephen Baker, Barney J Isaac, Ayyan Raj Neeravi, Dhiviya Prabaa Muthuirulandi Sethuvel, Biju George, Balaji Veeraraghavan

## Abstract

Hypervirulent *K. pneumoniae* (HvKp) is typically associated with ST23 clone; however, hvKp is also emerging from clones ST11, ST15 and ST147, which are also multi-drug resistant (MDR). Here, we aimed to characterise nine novel MDR hvKp isolates harbouring mosaic plasmids simultaneously carrying antimicrobial resistance (AMR) and virulence genes. Nine HvKp isolates obtained from hospitalised patients in southern India were characterized for antimicrobial susceptibility and hypervirulence phenotypes. All nine hvKp isolates were subjected to whole genome sequencing (WGS) using Ilumina HiSeq2500 and a subset of four were sequenced using Oxford Nanopore MinION. Among the nine isolates, seven were carbapenem-resistant, two of which carried *bla*_NDM-5_ on an IncFII plasmid and five carried *bla*_OXA-232_ on a ColKP3 plasmid. The virulence determinants were encoded in a mosaic plasmid (∼320 Kbp) that formed as a result of its insertion in a IncFIB-IncHI1B plasmid co-integrate. The mosaic plasmid carried AMR genes (*aadA2, armA, blaOXA-1, msrE, mphE, sul1* and *dfrA14*) in addition to *rmpA2, iutA* and *iucABCD* virulence genes. Interestingly the mosaic plasmid carried its own type IV-A3 CRISPR-cas system that is likely able to target the acquisition of IncF plasmid with the help of a *traL* spacer. The convergence of virulence and AMR is the biggest threat among invasive *K. pneumoniae* infections. However, increasing reports of the presence of mosaic plasmid carrying both AMR and virulence genes suggests MDR-hvKp isolates are no longer confined to selected clones and the containment of such isolates is very challenging.

**IMPORTANCE:** *Klebsiella pneumoniae* is an opportunistic pathogen that commonly associated with hospital-acquired infections in the urinary tract, respiratory tract, lung, wound sites. The organism has gained notoriety by acquiring additional genetic traits to become either hypervirulent (HV) phenotype or multidrug resistant (MDR) phenotype. Though the infections by both these phenotypes were very challenging to treat, the MDR *K. pneumonia* (MDR-Kp) were remained in the hospital settings while HV *K. pneumonia* (hvKp) strains were mostly originated from the community settings. In a recent turn of events, the evolution of MDR-Kp and hvKp has converged as both clones found to carry both MDR plasmids and virulence plasmid. These convergent strains are challenging to treat and is associated with higher mortality rate. As the recent hvKp isolates harbour mosaic plasmid encoding both AMR and virulence determinants there is a need to investigate the evolution of these pathogens. The significance of our research is in characterising the novel mosaic plasmid identified in MDR-hvKp isolates that belong sequence type (ST) 2096. Tracking the possible evolution pathway of MDR-hvKPs would greatly help in the proper surveillance and management of this superbugs.

**Repositories:** The whole genome sequences of the present study isolates have been deposited in GenBank, NCBI, with accession numbers CP053765 - CP053770, CP053771 – CP053780, CP058798-CP058806, JAARNO010000001.1 - JAARNO010000005.1, JAAQSG000000000, JAARNJ000000000, JAARMH000000000 and JAAQTC000000000

## INTRODUCTION

Hypervirulent *K. pneumoniae* (HvKp) is becoming more prevalent globally and being associated with an increasing number of fatalities (1, 2). Early reports of hvKp highlighted an association with liver abscesses, but more recent reports have documented infections in various internal sites in patients without liver abscesses (3,4). HvKp isolates were documented to be limited in their antimicrobial resistance (AMR) phenotypes because of a barrier for acquiring MDR plasmids (5). However, there has been the emergence of multidrug resistant (MDR) variants more recently (6). The acquisition of mobile genetic elements harbouring carbapenemases among hvKp isolates, or the uptake of pLVPK-like virulence plasmid in carbapenem resistant *K. pneumoniae* (CRKP) results in the dangerous convergence of carbapenem-resistant and hypervirulent phenotypes (7).

The population structure of hvKp, when assessed by multi-locus sequence typing (MLST) and whole-genome sequencing (WGS), indicates that most hvKp isolates belong to clonal groups (CG) 23, 65, 86, 375, and 380 (8). Conversely, CRKp is associated with a clonal expansion of CG258 in Europe, while its endemic dissemination in Asia is associated with ST11, ST14, ST147, ST149, and ST231 (9). The convergence of AMR and virulence, irrespective of the clonal group, has transformed *K. pneumoniae* into a group pathogen that cause serious infections with very limited treatment options (10).

Convergent strains of carbapenem-resistant hypervirulent *K. pneumoniae* (CR-hvKp) have undergone frequent genetic transposition through the formation of a fusion/co-integrate or mosaic plasmid (11). Notably, these mosaic plasmid are typically composed of two different plasmid backbones and generate the potential for AMR and virulence determinants to be encoded within a single plasmid (12). Similar mosaic plasmid with diverse backbones carrying IncFIB-IncHI1B, IncFIBK-IncHI1B, IncFIB-IncR have been reported in China (11, 13, 14). These emerging HvKp are extremely concerning as these superbugs have the potential to cause devastating hospital outbreaks (15, 16). Understanding of the diversity of mosaic plasmid held within CR-hvKp isolates is currently limited due to a lack of completely assembled circular plasmids. Here, we characterised nine MDR ST2096 hvKp carrying mosaic plasmid that simultaneously encoded both antimicrobial resistance and virulence genes. The complete genome sequence of a subset of four isolates were further characterised aiming to elucidate the structure of mosaic plasmid through comparative genomics.

## MATERIALS AND METHODS

### Bacterial isolates

The nine studied *K. pneumoniae* were isolated from patients with bacteraemia admitted to Christian Medical College, Vellore, India in 2019. The isolates were identified using standard biochemical methods and further confirmed by Vitek-MS (Database v2.0, bioMerieux, France). The isolates were screened for a hypermucoviscous phenotype using the string test (2). In addition, the mucoid phenotypic genes *rmpA* and *rmpA2* were screened using PCR, as described previously (17,18). The demographic and clinical details of the nine patients were collected from electronic medical records maintained at the Hospital. The study was approved by Institutional Review Board of Christian Medical College, Vellore with minute number 9,616 (01/09/2015).

### Antimicrobial susceptibility testing

Antimicrobial susceptibility testing (AST) was performed by Kirby Bauer disc diffusion method according to CLSI 2019 guidelines (19). The tested antimicrobials were, cefotaxime (30µg), ceftazidime (30µg), piperacillin/tazobactam (100/10µg), cefoperazone/sulbactam (75/30µg), imipenem (10µg), meropenem (10µg), ciprofloxacin (5µg), levofloxacin (5µg), amikacin (30µg), gentamicin (10µg) and minocycline (30µg). The minimum inhibitory concentration (MIC) against Meropenem was determined by broth micro dilution (BMD). *Escherichia coli* ATCC 25922, *Enterococcus faecium* ATCC 29212 and *Pseudomonas aeruginosa* ATCC 27853 were used as the quality control strains for AST. AST results were interpreted according to CLSI guidelines (20).

### DNA extraction and genome sequencing

The study isolates were revived from the archive at Department of Clinical Microbiology and single isolated colony was grown in LB broth (Oxoid, Hampshire, United Kingdom) at 37°C. Total genomic DNA was extracted from the pelleted cells using Wizard DNA purification kit (Promega, WI, USA) as per the manufacturer’s protocol. Extracted DNA was quantified using NanoDrop One spectrophotometry (Thermo Fisher Scientific, MA, USA) and Qubit 3.0 fluorometry (Life Technologies, CA, USA) and stored at −20^°^C until further use.

Sequencing library was prepared using the Nextra DNA Flex library preparation kit (Illumina, San Diego, CA). as per the manufacturer’s instructions. Subsequently the paired end library was subjected to sequencing on a HiSeq 2500 platform (Illumina, USA) generating 2 x 150-bp reads. Sequencing reads with a PHRED quality score below 20 were discarded and adapters were trimmed using cutadapt v1.8.1 and assessed with FastQC v0.11.4. For a subset of four isolates, long read sequencing was carried out using Oxford Nanopore MinION platform with FLO-MIN106 R9 flow cell (Oxford Nanopore Technologies, Oxford, UK). Long read DNA library was prepared using the SQK-LSK108 ligation sequencing kit (v.R9) along with ONT EXP-NBD103 Native Barcode Expansion kit following the manufacturer’s protocol (Oxford Nanopore Technologies, Oxford, UK). The library was loaded onto the flow cells, run for 48 hrs using the standard MinKNOW software. The Fast5 files generated from MinION sequencing were subjected to base calling using Guppy (https://github.com/gnetsanet/ONT-GUPPY).

### Genome assembly and evaluation

Draft genome sequence data generated using Illumina were assembled using SPAdes (v.3.13.0) (21). For a subset of four isolates complete and highly accurate assembly was achieved using hybrid *de novo* assembly approach (22). The nanopore long reads were error-corrected with the standalone Canu error correction tool (v.1.7) and assembled using the Unicycler hybrid assembly pipeline (v 0.4.6) with the default settings (23, 24). The obtained genome sequence was polished using high quality Illumina reads as described previously (25). The assembled complete genome was subjected to quality assessment using CheckM v1.0.5 (26) and Quast v4.5 (27). CheckM estimated the completeness and contiguity while Quast was used to detect mis-assemblies, mismatches and indels by aligning the assemblies with the reference genome, *K. pneumoniae* NTUH-K2044 (AP006725). Since *K. pneumoniae* NTUH-K2044 is a well characterised-type strain of ST23 hypervirulent *K. pneumoniae*, it was used as the reference genome.

### Genome analysis

Genome assemblies were submitted to NCBI GenBank and annotated using the Prokaryotic Genome Annotation Pipeline (PGAP v.4.1) from NCBI (28). The resistance profile of the assembled genome sequences was Resfinder 4.1 available from CGE server (https://cge.cbs.dtu.dk/services/ResFinder/). Similarly, the presence of plasmids in the genomes was identified and characterized using PlasmidFinder (v.1.3) available at CGE server (https://cge.cbs.dtu.dk/services/PlasmidFinder). Further, MLST and virulence locus (yersiniabactin, aerobactin and other siderophore production systems) were identified using Kleborate (v.2.0.0) (https://github.com/katholt/Kleborate) (29). The presence of virulence factors were confirmed using virulence database at Pasteur Institute for *K. pneumoniae* (https://bigsdb.pasteur.fr/cgibin/bigsdb/bigsdb.pl?db=pubmlst_klebsiella_seqdef_public&page=sequenceQuery). The K and O antigen loci were also identified using Kaptive available at Kleborate (30). The final assembled circular chromosome and plasmid were visualized using CGview server v.1.0 and Easyfig (32). CRISPR regions in the genomes were identified with CRISPRCasTyper web server (http://cctyper.crispr.dk) (33). The genetic distance between the isolates were calculated using average nucleotide identity (ANI) available at OrthoANI (34). Pairwise distance between the nine isolates were determined with BA10835 as reference using SNP-dists v 0.6.3 (34) from the raw reads (https://github.com/tseemann/snp-dists) as described previously (35).

## RESULTS

### Clinical manifestations and microbiological characteristics of isolates

The demographic and clinical details of the patients with hvKp bacteraemia are shown in Table 1. The nine selected *K. pneumoniae* isolates were resistant to all tested antimicrobials by disc diffusion and were initially considered to be extensively drug resistant (XDR). However, on determining MIC, one of the isolates was found to be susceptible to meropenem (MIC ≤0.5µg/ml). All organisms generated a negative string test, but were positive for the *rmpA2* gene by PCR. Based on MLST, all nine *K. pneumoniae* isolates belonged to ST2096, a single locus variant of ST14. The surface capsule (K) loci were predicted to be K64 and the O-antigen encoding loci was determine to be O1v1 in all isolates.

**Table 1.**
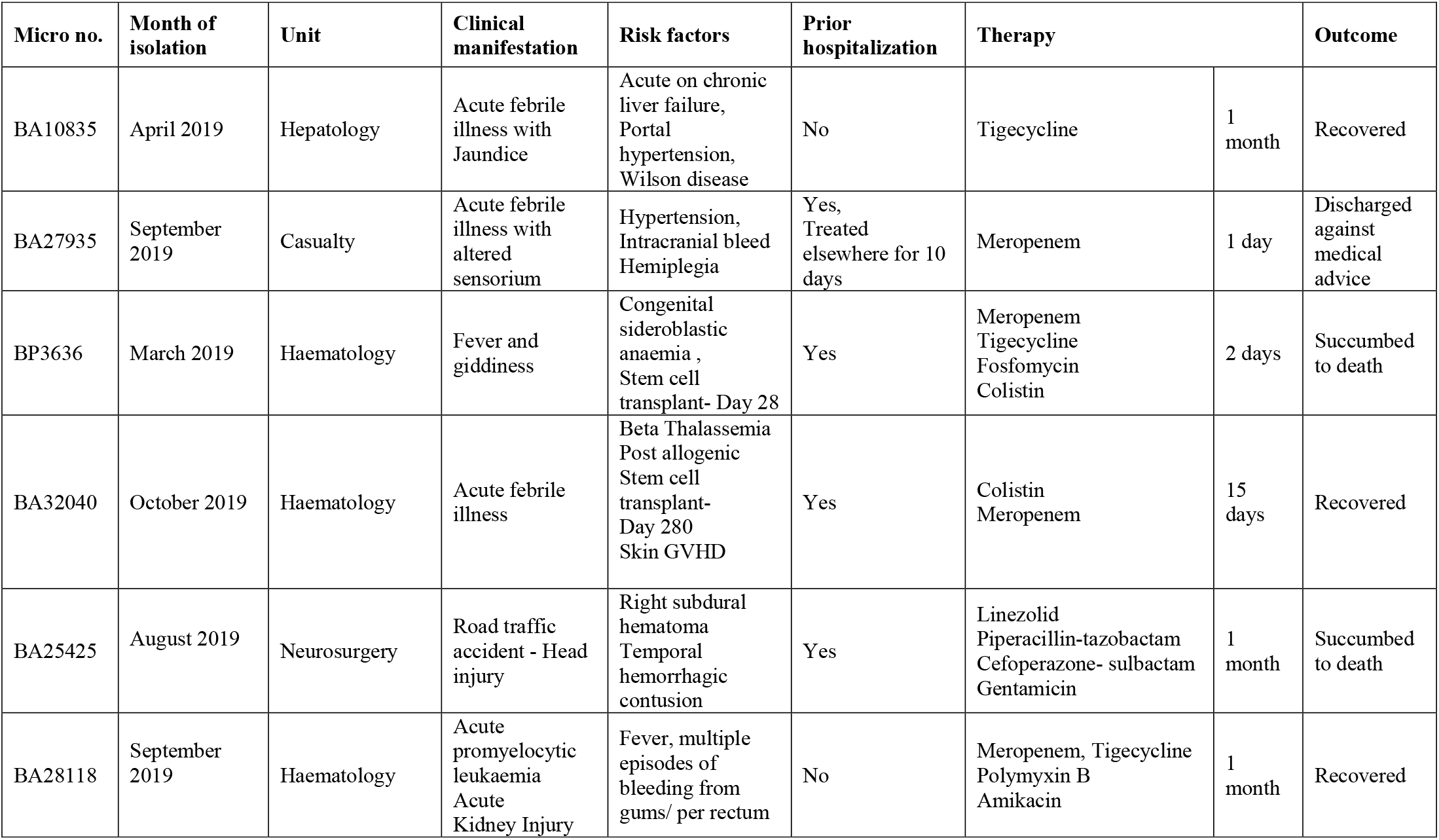
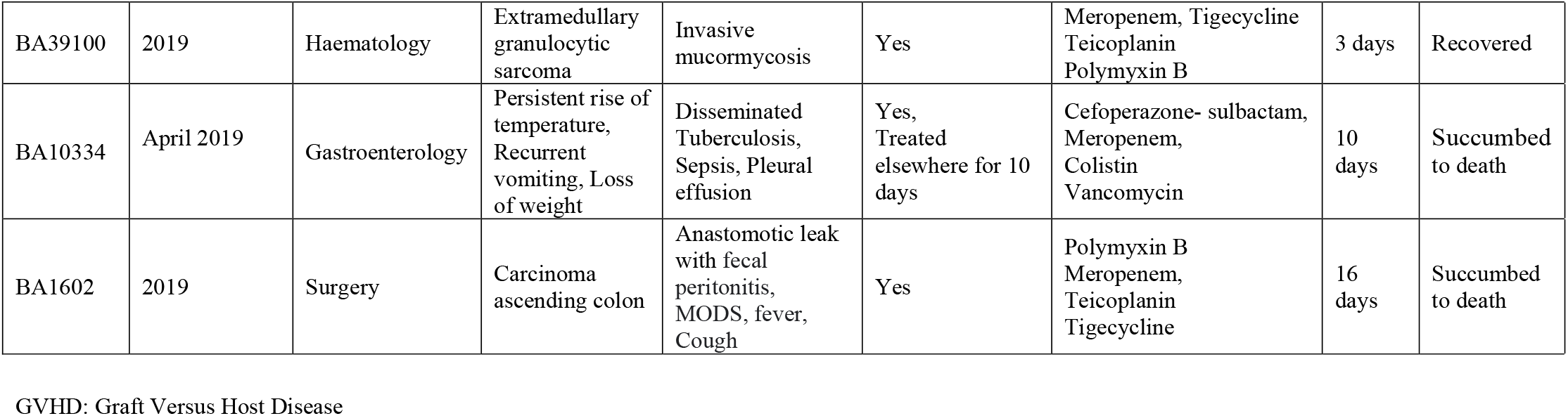
The demographic and clinical details of the four patients with hypervirulent *K. pneumoniae* bacteraemia

The pairwise average nucleotide identity (ANI) between the nine draft genomes showed >99.80% similarity (Suppl Fig. 1a). Pairwise SNP difference between the nine study isolates identified the two clusters of strains with isolates BA10835 and BA27935 being >260 SNP distant from the remaining seven (Suppl Fig 1b). Among the cluster containing 7 isolates, strain BA10334 and BA1602 were closely related (2 SNPs) so does strain BA25425 and BP3636. Our analysis was successful in identifying a possible two outbreaks of ST2096 in the hospital settings though more details are required to confirm our hypothesis.

### Antimicrobial resistance determinants

The AMR determinants of four complete genomes are listed in Table 2 and five draft genomes are listed is Table 3. The nine hvKp isolates were found to possess an array of AMR genes associated with multiple plasmids (Table 2 and Table 3). Atypically, we found that 3/4 isolates with complete genomes had *aac(6’)-lb-cr, bla*_OXA-1_ and *dfrA1*, integrated into chromosome on mobile genetic elements. Specifically, *aac(6’)-lb-cr* and *bla*_OXA-1_ were associated with an IS26 and were inserted in the middle of the chromosome at position ∼2.3Mbp, while *dfrA1* was associated with an IS*Kpn26* and class 1 integron and was inserted at position ∼5.3Mbp. Additionally, a duplicate region of 7bp (AGTCCGT) flanked the AMR genes where IS26 was inserted (**Figure 1**). Two isolates (BA10835 and BA27935) were also found to carry *bla*_NDM-5_ on IncFII plasmid and one isolate (BP3636) carried *bla*_OXA-232_ on ColKp3 plasmid. The fourth isolate, BA32040, which was susceptible to meropenem, lacked a carbapenemase encoding gene.

**Figure 1:**
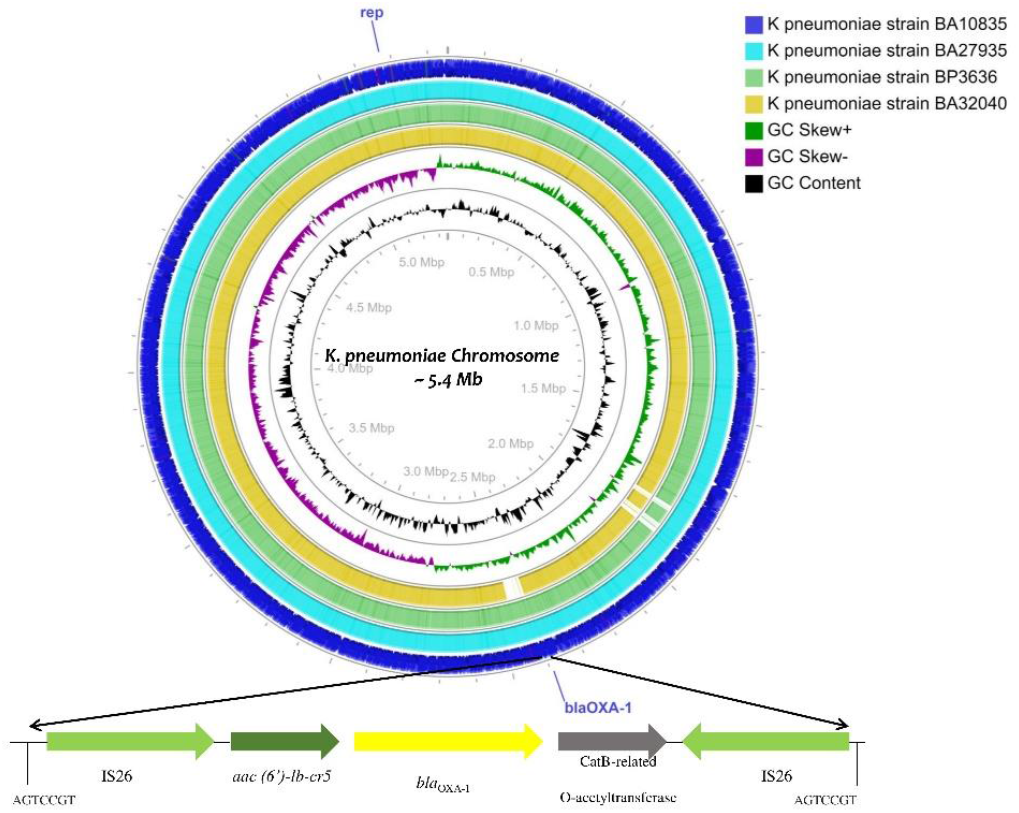
Circular genome maps of four ST2096 MDR hypervirulent *K. pneumoniae* chromosomes generated using CGview server (Grant and Stothard, 2008). Circles from the outside to the inside show the CDS region of strain BA10835 (blue), strain BA27935 (Cyan), BP3636 (Green), BA32040 (Yellow), GC skew (dark green and Magenta), GC content (black). Linear view of the IS26 mediated translocatable units carrying *aac(6′)-Ib-cr* (fluoroquinolones and aminoglycosides), *bla*_OXA–1_ (ampicillin), catB3 (chloramphenicol) inserted to the chromosome. A repeat region of 7 bases read as AGTCCGT was present on either ends where the insertion was observed.

**Table 2:** Phenotypic and genotypic characteristics obtained using hybrid genome assembly, of four ST2096 Indian MDR hypervirulent *K. pneumoniae* in comparison with the previously reported isolates with hybrid plasmid

| Isolate ID | BA10835 | BA27935 | BP3636 | BA32040 | Lam et al., 2019 | Turton et al., 2019 |  |
| --- | --- | --- | --- | --- | --- | --- | --- |
| MLST | ST2096 | ST2096 | ST2096 | ST2096 | ST15 | ST147* | ST383* |
| Accession numbers | CP053765 - CP053770 | CP058798-CP058806 | CP053771 – CP053780 | JAARNO010000001.1 to JAARNO010000005.1 | ERR2866472<br>ERR2866473 | NZ_MZMY01000000<br>CP040724.1 | CP034200.2 |
| Meropenem MIC | 128 µg/ml | 4 µg/ml | 64 µg/ml | ≤0.5µg/ml | NA | NA | NA |
| Chromosomal AMR genes | *aac(6')-lb-cr*, *bla<sub>SHV</sub>*, *bla<sub>OXA-1</sub>*, *fosA*, *dfrA1* | *aac(6')-lb-cr*, *bla<sub>OXA-1</sub>*, *bla<sub>SHV</sub>*, *fosA*, *dfrA1* | *bla<sub>SHV</sub>*, *fosA*, *dfrA1* | *aac(6')-lb-cr*, *bla<sub>SHV</sub>*, *bla<sub>OXA-1</sub>*, *fosA*, *dfrA1* | *bla<sub>SHV-5</sub>*, *fosA*, *dfrA1* | *bla<sub>SHV-67</sub>*, *fosA*, *dfrA1* | *bla<sub>SHV-26</sub>*, *fosA*, *dfrA1* |
| Chromosomal virulence genes | *fyuA*, *irp1*, *kfuABC*, *mrkACFJ*, *ybtAEPQSTUX* | *fyuA*, *irp1*, *kfuABC*, *mrkACFJ*, *ybtAEPQSTUX* | *fyuA*, *irp1*, *irp2*, *kfuABC*, *mrkABCFHJI*, *ybtAEPQSTUX* | *fyuA*, *irp1*, *kfuABC*, *mrkABCFHJI*, *ybtAEPQSTUX* | *fyuA*, *irp1*, *kfuABC*, *mrkABCFHJI*, *ybtAEPQSTUX* | *fyuA*, *irp1*, *irp2*, *mrkABCFHJI*, *ybtAEPQSTUX* | *mrkABCFHJI*, *pld1* |
| No. of plasmids | 5 | 4 | 4 | 4 | 4 | 3 | 2 |
| IncHI1B /IncFIB (pNDM-MAR) | *aadA2*, *armA*, *bla<sub>TEM-1B</sub>*, *bla<sub>CTX-M-15</sub>*, *mphE*, *msrE*, *sul1*, *tetD*, *dfrA12* | *aadA2*, *armA*, *bla<sub>TEM-1</sub>*, *bla<sub>CTX-M-15</sub>*, *mphE*, *msrE*, *tetD*, *dfrA12*, *sul1* | *aac(6')-lb-cr*, *aadA2*, *armA*, *bla<sub>TEM-1A</sub>*, *bla<sub>CTX-M-15</sub>*, *bla<sub>OXA-1</sub>*, *msrE*, *mphE*, *sul1*, *tetD*, *dfrA12*, *dfrA14* | *bla<sub>TEM-1A</sub>*, *bla<sub>CTX-M-15</sub>*, *tetD*, *dfrA14* | *Aac3'-IIa*, *aadA1*, *bla<sub>TEM</sub>*, *bla<sub>CTX-M-15</sub>*, *sul1*, *dfrA1*, *sat2* | *sul1*, *sul2*, *armA*, *dfrA5*, *mph(A)*, *msr(E)*, *mph(E)*, *aph(3')-Ia* | *bla<sub>NDM-5</sub>*, *bla<sub>CTX-M-15</sub>*, *bla<sub>OXA-9</sub>*, *qnrS1*, *bla<sub>TEM-1B</sub>*, *dfrA5*, *catA1*, *sul1*, *sul2*, *armA*, *aph(3)-Ia*, *aph(3)-VI*, *aac(6)-lb*, *aadA1*, *aac(6)-lb-cr*, *mph(A)*, *mph(E)*, *msr(E)* |
|  | *iucABCD*, *iutA*, *rmpA2* | *iucABCD*, *iutA*, *rmpA2* | *iucABCD*, *iutA*, *rmpA2* | *iucABCD*, *iutA*, *rmpA2* | *iucABCD*, *rmpA2* | *iutA*, *iucABCD*, *rmpA*, *rmpA2*, *terABCDEWXYZ*, *cobW*, *luxR*, *pagO*, *shiF* | *utA*, *iucABCD*, *rmpA/rmpA2*, *terABCDEWXYZ*, *cobW*, *luxR*, *pagO*, *shiF* |
| IncFIBK | *catA1* | absent | No AMR gene | No AMR gene | Absent | absent | absent |
| ColKP3 | absent | absent | *bla<sub>OXA-232</sub>* | absent | Absent | absent | absent |
| IncFII | *aadA2*, *rmtB*, *bla<sub>NDM-5</sub>*, *ermB*, *mphA*, *sul1*, *dfrA12* | *aadA2*, *rmtB*, *bla<sub>NDM-5</sub>*, *bla<sub>TEM-1</sub>*, *ermB*, *mphA*, *sul1*, *dfrA12* | absent | *rmtB*, *bla<sub>TEM-1B</sub>*, *ermB*, *mphA* | *aacA4*, *bla<sub>OXA-1</sub>*, *bla<sub>TEM-1</sub>*, *cat* | absent | absent |
| Other plasmids | ColRNAI | Col(BS512), ColRNAI | ColRNAI | ColRNAI | Col4401, ColpVC | IncFIB (pQil) | absent |

MIC: Minimum Inhibitory Concentration determined by broth micro dilution; AMR: antimicrobial resistance; \*Virulence plasmid was a hybrid of IncFII(K)/IncFIB(K)

**Table 3:** Genotypic characteristics of multidrug resistant hypervirulent *K. pneumoniae* belonging to ST2096 obtained from short read assembly.

| Accession number | *rmpA*<br>and/ or<br>*rmpA2* | Capsule<br>type | O<br>antigen | Ybt,<br>ICEKp | Resistance genes | Plasmids | Virulence genes |
| --- | --- | --- | --- | --- | --- | --- | --- |
| JAARMH000000000<br>BA1602 | *rmpA2* * | K64 | O1v1 | *ybt14*;<br>ICEKp5 | *aac(6)-lb-cr*, *aadA2*, *armA*, *blaCTX-M-15*, *blaOXA-1*, *blaSHV-106*, *blaTEM-150*, *dfrA1*, *dfrA12*, *dfrA14*, *fosA6*, *mphE*, *msrE*, *sul1*, *tetD* | ColKP3, IncFIBK, incFIB (pNDM-MAR), IncHI1B (pNDM-MAR) | *fyuA*, *irp1*, *irp2*, *kfuAB*, aerobactin, *mrkABCDHFIIJ* |
| JAAQTC000000000<br>BA25425 | *rmpA2* * | K64 | O1v1 | *ybt14*;<br>ICEKp5 | *aadA2*, *ArmA*, *Sat-2A*, *blaOXA-1*, *blaSHV-28*, *blaTEM-1D*, *blaOXA-232*, *blaCTX-M-15*, *mphE*, *msrE*, *sul1*, *tetD*, *dfrA1*, *dfrA12*, *dfrA14* | ColKP3, ColRNAI, IncFIBK, IncFIB (pNDM-Mar), IncHI1B (pNDM-MAR) | *fyuA*, *irp1*, *irp2*, *kfuABC*, *mrkABCDHFIIJ* |
| JAAQSG000000000<br>BA28118 | *rmpA2* * | K64 | O1v1 | *ybt14*;<br>ICEKp5 | *AadA2*, *ArmA*, *blaOXA-1*, *blaSHV-28*, *blaTEM-1D*, *blaOXA-232*, *blaCTX-M-15*, *mphE*, *msrE*, *sul1*, *tetD*, *dfrA12*, *dfrA14* | IncFIBK, IncFIB(pNDM-Mar), ColKP3, ColBS512, IncHI1B (pNDM-MAR) | *fyuA*, *irp1*, *irp2*, aerobactin, *kfuABC*, *mrkABCDHFIIJ* |
| JAARNJ000000000<br>BA39100 | *rmpA2* | K64 | O1v1 | *ybt14*;<br>ICEKp5 | *aac(6)-lb-cr*, *aadA2*, *armA*, *sat2A*, *blaCTX-M-15*, *blaOXA-1*, *blaOXA-232*, *blaSHV-106*, *blaTEM-1*, *catB*, *dfrA1*, *dfrA12*, *dfrA14*, *fosA6*, *mphE*, *msrE*, *sul1*, *tetD* | ColKP3, IncFIBK, IncFIB(pNDM-Mar), IncHI1B (pNDM-MAR) | *fyuA*, *irp1*, *irp2*, *kfuA*, *kfuC*, aerobactin, *mrkABCDHFIIJ* |
| JAAQSS000000000<br>BA10334 | *rmpA2* * | K64 | O1v1 | *ybt14*;<br>ICEKp5 | *aac(6)-lb-cr*, *aadA2*, *armA*, *blaCTX-M-15*, *blaOXA-1*, *blaOXA-232*, *blaSHV*, *blaTEM-1A*, *fosA*, *msrE*, *mphE*, *sul1*, *tetD*, *dfrA1*, *dfrA12*, *dfrA14* | ColKP3, IncHI1B (pNDM-MAR), IncFIB (pNDM_MAR), IncFIBK | *fyuA*, *irp1*, *irp2*, *iutA*, aerobactin, *kfuABC*, *mrkABCDHFIIJ* |

*rmpA2*\*: frameshift mutation in *rmpA2*

### Plasmids

All nine isolates each carried either four or five plasmids including a virulence plasmid. The plasmids and their associated ARGs as deduced by complete genomes of four isolates are shown in Table 2. Notably, *bla*_NDM-5_ was carried on IncFII plasmid (∼97Kbp) along with additional ARGs such as *aadA2, rmtB, ermB, mphA, sul1, dfrA12* and *bla*_TEM-1_ (**Figure 2**). We also found a segment of IS30 family transposase of 293bp, with identity similar to IS*Aba125* adjacent to *bla*_NDM-5_. The closest matching plasmid from a global database was from *K. pneumoniae* JUNP 055 (LC506718), which also harboured *bla*_NDM-5_. This plasmid (LC506718) shared ∼80% sequence identity to that of *E. coli* M105 (AP018136), which lacked *bla*_NDM-5_.

**Figure 2:**
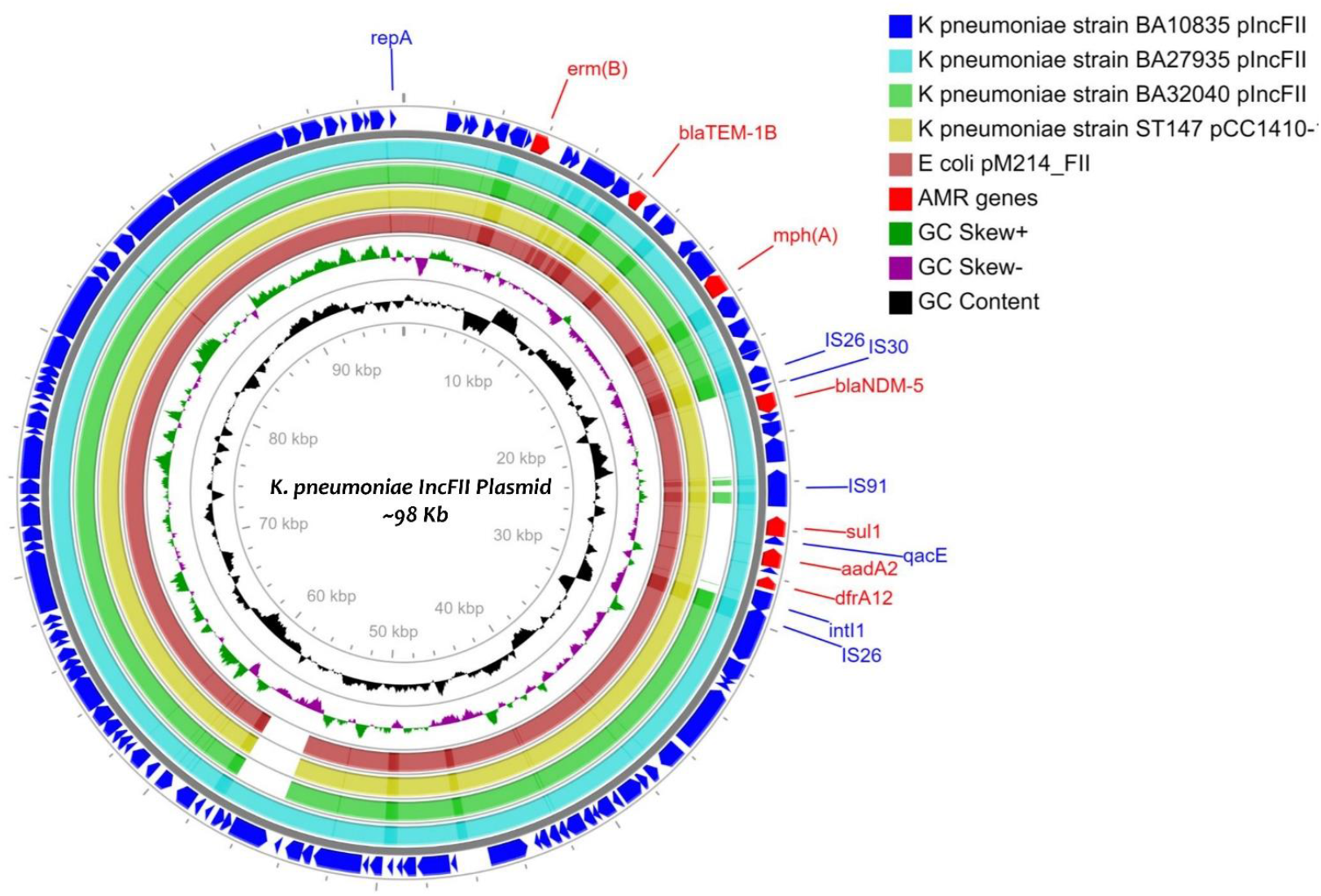
Circular genome maps of IncFII plasmids of three MDR hypervirulent *K. pneumoniae* belonging to ST2096 generated using CGview server. Circles from the outside to the inside show the CDS region of strain BA10835 (blue), strain BA27935 (Cyan) and BA32040 (Green) and nearest matching reference plasmids that belong to *K. pneumoniae* pCC1410-1 (Yellow; KT725788) and *E. coli* pM214 (Red; AP018144).GC skew (dark green and Magenta), GC content (black) of the plasmid are represented in the inner circles. IncFII carries several antimicrobial resistance genes including *bla*_NDM-5_.

An additional large (∼307Kbp) plasmid was present in all four genomes subjected to hybrid assembly and was a fusion of IncFIB and IncHI1B backbones and carried both ARGs and HvKp virulence genes (Table2). However, this plasmid was found to be shorter in BA32040 (272Kbp) and lacked the *aadA2, armA, bla*_OXA-1_, *msrE, mphE, sul1* and *dfrA14* ARGs in comparison to plasmids in the other three HvKp. Two plasmids in the study isolates, Col (BS512) and ColRNAI, did not possess any ARGs. Our data suggest that all the nine HvKp harboured a virulence plasmid which was a fusion of IncHI1B and IncFIB (Table 2 and Table 3).

### Virulence

The key virulence determinant carried by the chromosome of *K. pneumoniae* is the *ybt* locus, which is mobilized by ICEKp. In all the nine isolates, *ybt14* was carried on ICEKp5 and integrated into the chromosome. Yersiniabactin receptors, such as *fyuA* and *irp1*, were also present on the chromosome, along with an alternative *kfu* gene cluster encoding for iron uptake and *mrk* gene cluster which facilities biofilm formation.

The presence of a large mosaic virulence plasmid of ∼307Kbp was the hallmark of these ST2096 isolates with key virulence determinants, such as *rmpA2* and aerobactin siderophore, which is encoded by *iucABCD* and *iutA* coding for its receptor. Notably, a frameshift mutation was observed in *rmpA2* among all the isolates which presumably was associated with a negative string test result. **Figure 3a** shows a comparison of mosaic plasmid from the present study with those previously reported *K. pneumoniae* belonging to diverse sequence types. As depicted in **Figure 3b**, the large mosaic plasmid from the study isolates were compared with IncFIB-IncHI1B co-integrate plasmid (CP006799 and AP018748) as well as the reference virulence plasmid (pLVPK: AY378100.1). In addition to the virulence genes, the mosaic virulence plasmid also carried genes encoding heavy metal resistance; *merARCTP* (mercury) and *terBEDWXZ* (tellurium).

**Figure 3a:**
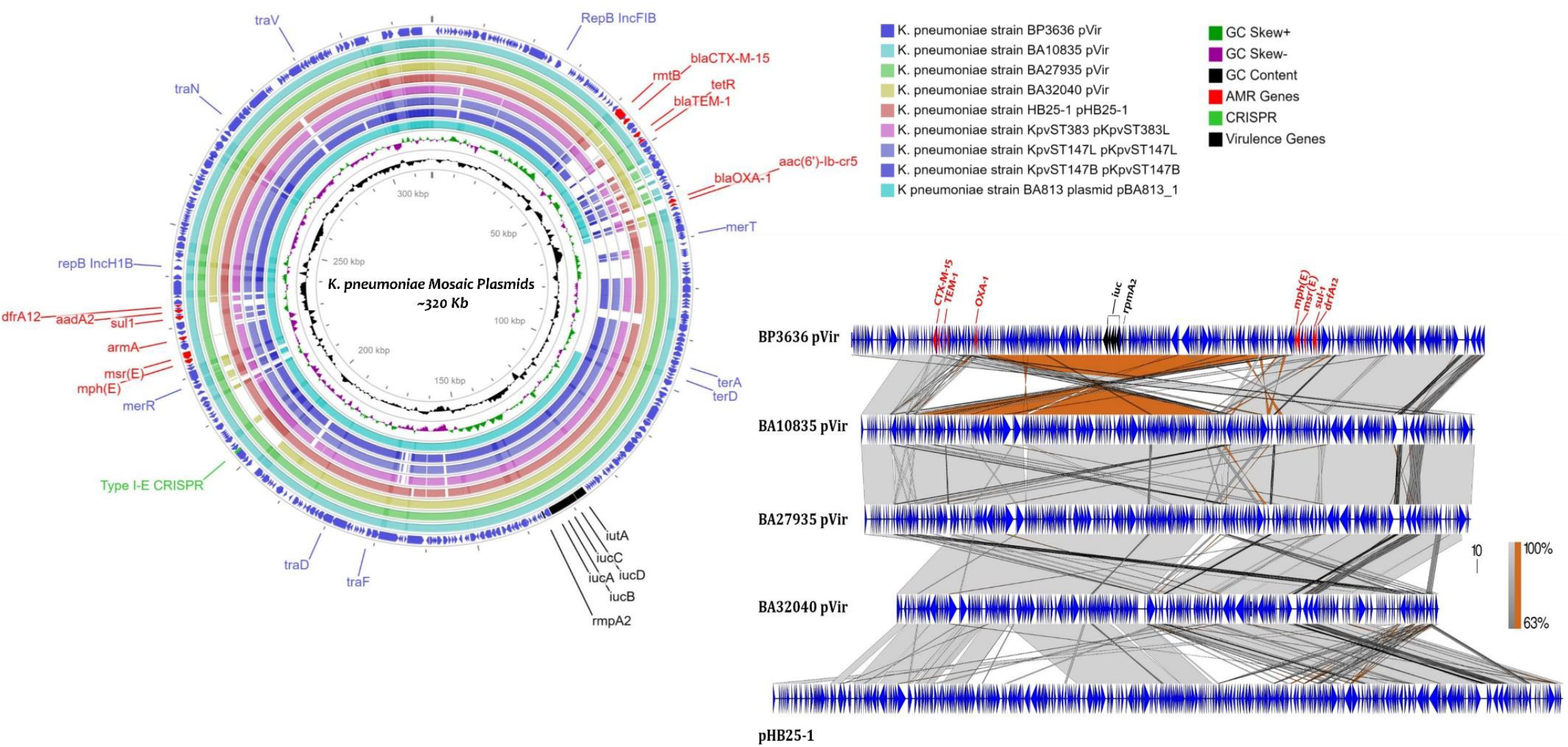
Circular genome maps of mosaic plasmid generated using CGview server depicting the comparison of IncFIB-IncHIB-pVir mosaic plasmids of strain BP3636 (Navy blue), strain BA10835 (Cyan), BA27935 (Green), BA32040 (Yellow) with previously reported mosaic plasmids pHB25-1 (CP039526; Red), pKpvST383L (CP034201; Pink), pKpvST147L (CM007852; Light blue), pKpvST147B (CP040726; blue), pBA813 (MK649825; turquoise) from hvKp. Linear alignment prepared using Easyfig shows the comparison of mosaic plasmids with pHB25-1.

**Figure 3b:**
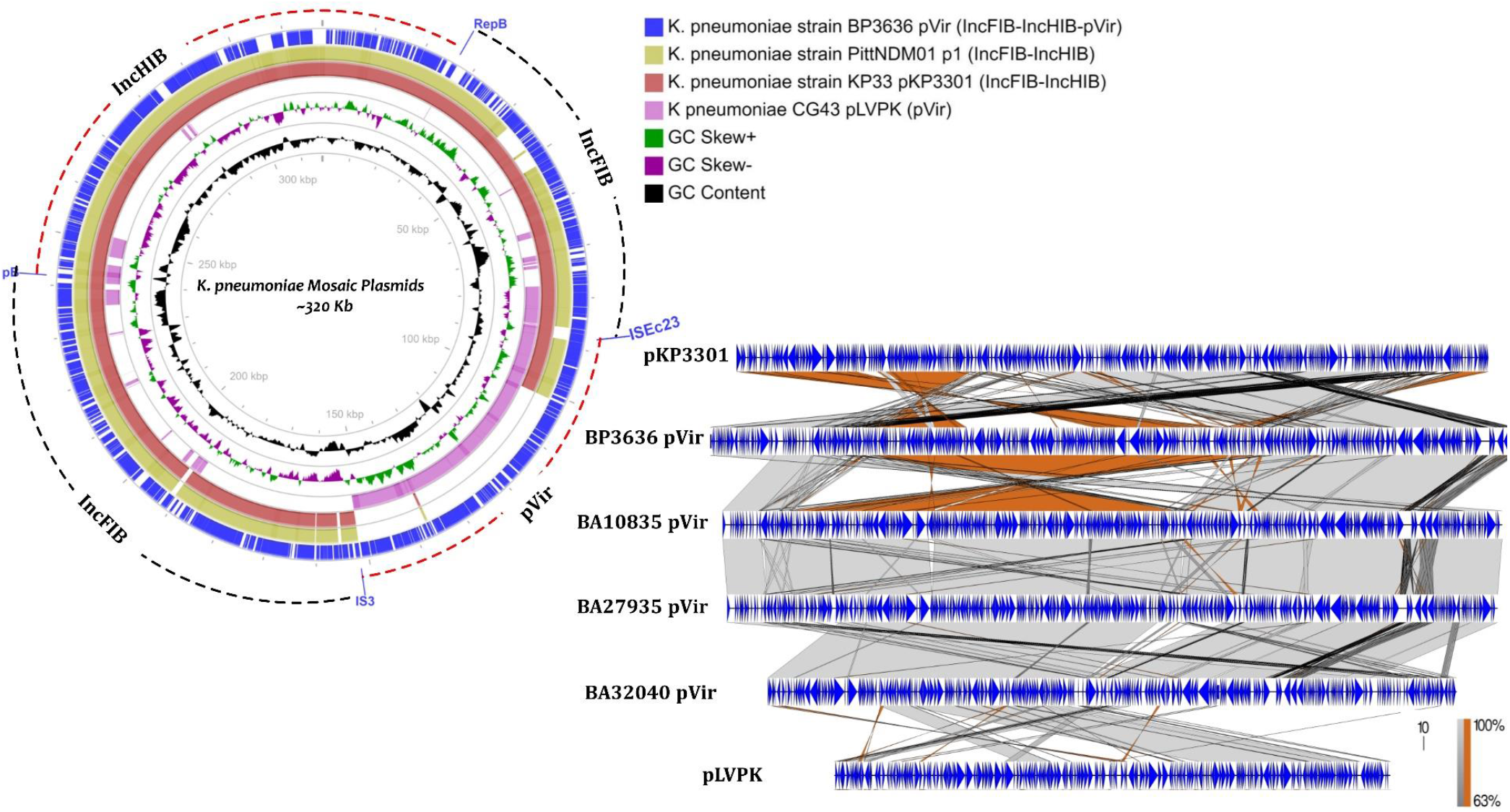
Circular genome maps generated using CGview server depicting the comparison of IncFIB-IncHIB-pVir mosaic plasmids of strain BP3636 (Navy blue) with IncFIB-IncHIB backbone of plasmid p1 (CP006799), pKP3301 (AP018748) and virulence plasmid pLVPK (NC_005249). Linear alignment prepared using Easyfig shows the possible insertion of the virulence associated region from a pLVPK like plasmid into the IncFIB-IncHIB backbone by means of transposases IS3and IS66.

### CRISPR-Cas system

From the nine-draft genome sequence, we found the occurrence of type I-E CRISPR and type IV-A3 CRISPR system in each genome (**Figure 4)**. Based on the information gathered from the four complete genome sequence, Type I-E CRISPR-Cas system was carried by the chromosome and type IV-A3 system is located on the mosaic plasmid. The number of spacers ranged from 7 to 12, which were 32bp in length. Adjacent to the CRISPR array was IS*Kpn26*. These plasmid CRISPR regions were characterized by the presence of 5-12 spacers and a 29bp repeat region. One spacer each from the mosaic plasmid of the three isolates was similar to *traL* of IncF plasmids found in *K. pneumoniae*.

**Figure 4:**
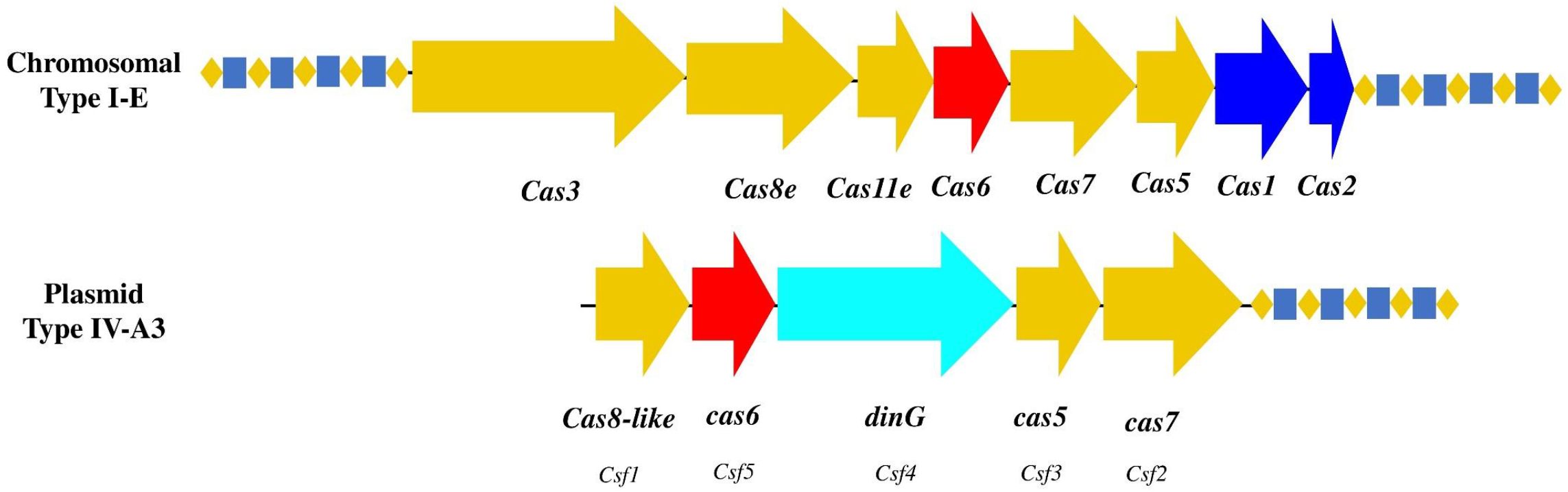
Schematic representation of the CRISPR-cas system associated with ST2096 MDR hypervirulent *K. pneumoniae* **a**. Gene organization of the chromosomal type I-E CRISPR-cas system. b. Gene organization of the plasmid associated type IV-A3 CRISPR-cas system. Genes are colour-coded and labelled according to the protein families as represented by CRISPRCasTyper web server.

## DISCUSSION

The evolution of *K. pneumoniae* clinical isolates by the acquisition of multiple resistance plasmids has placed this species among the most important causative agent of nosocomial infections. These highly challenging MDR-Kp or CR-Kp isolates is majorly associated with the NDM or/and OXA-48-like carrying ST11, ST14, ST15, ST101, ST147, ST395 and ST231 in Asia and KPC harbouring ST258 in North America and Europe (36). Simultaneously, the classical *K. pneumoniae* (CKp) has acquired pLVPK-like virulence plasmid to become the hvKp that induces the community-acquired invasive infections (37). The hypervirulent phenotypes were mainly clustered into the clonal group CG23, which includes the sequence types ST23, ST26, ST57, and ST163 (38). This bidirectional convergence of divergently evolved populations resulted in the emergence of MDR-hvKp/CR-hvKp isolates within the nosocomial clones. The outbreak of nosocomial clones carrying virulence plasmid is a matter of major public health concern (39-41).

Among the nosocomial MDR clones, at present ST11 (15, 39), ST14 (42), ST15 (10, 43), ST231 (44), ST36 (45), ST437 (46) and ST2096 (44, 47) were reported to have acquired the pLVPK-like virulence plasmid. Rapid acquisition of virulence plasmid by these nosocomial clones in the last few years suggests these clones are now ready for nosocomial and health care associated outbreaks (15, 39). The present study data also support the concept of the increasing number of nosocomial infections due to the convergent hvKp clones (Table 1). Such unexpected emergence of MDR-hvKp ST2096 carrying ARGs and virulence genes in a fusion plasmid is extremely problematic (15).

The recent reports of the independent emergence of convergent hvKp isolates with mosaic plasmid in multiple geographical locations has made these organisms the latest superbug (2). The complete genome sequence of four study isolates resolved the plasmid structure and identified four plasmids each among the isolates except for strain BA10835 that carried five plasmids (Table 2). The ∼30 kb mosaic plasmid carrying both virulence and resistance determinants were comparable to previously reported fusion plasmids CM007852 CP034201, CP040726 (12) and MK649825 (44). Remarkably, these reference plasmids except MK649825 were from a diverse collection of clones (i.e. ST147 and ST383), which were found to harbour NDM and OXA-48 resistance genes. Moreover, the insertion of resistance cassette carrying ARGs such as *aadA2, armA, bla*_TEM-1B_, *bla*_CTX-M-15_, *mphE, msrE, sul1* and *dfrA12* of mosaic plasmid of independent origin are of serious concern. Interestingly the mosaic plasmid of Indian origin found to be fusion of IncFIB_K_/ IncHI1B backbone while the reference plasmids were emerged as a result of the fusion process of IncFII_K_ and IncFIB_K_ backbones (10, 12). The similarity among the plasmids with different backbones was attributed to the complement ARGs, MGEs and virulence genes encoded by them. Among these hvKp that harbours the fusion plasmids, mosaic structures were formed by the likely integration of virulence region from the hv plasmid into IncF-IncH co-integrate resistance plasmid (48). Diagrammatic representation of the possible evolution pathway of MDR-hvKP harbouring mosaic plasmid is represented in **Figure 5**.

**Figure 5:**
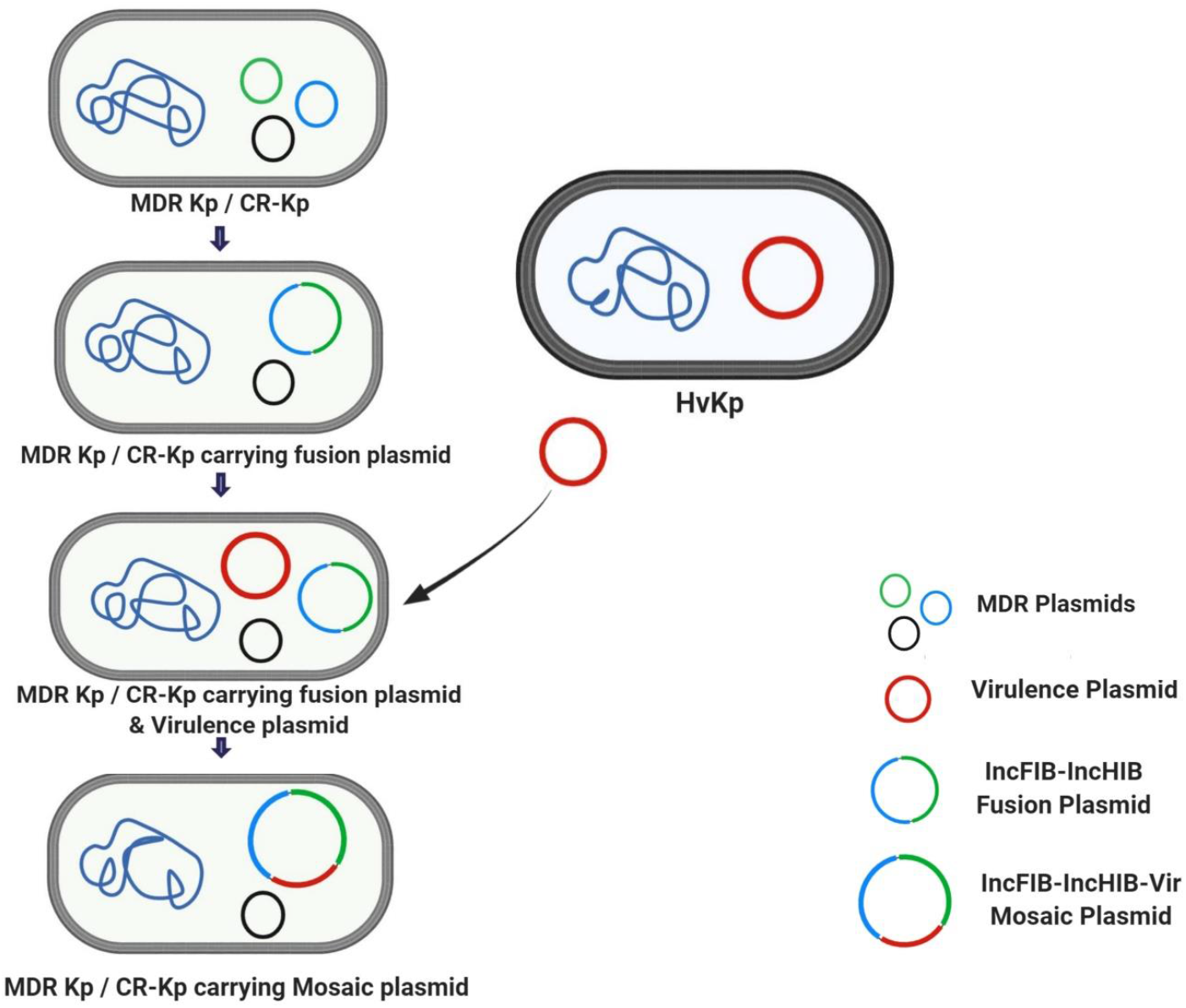
Schematic representation of the possible evolution pathway of ST2096 MDR hypervirulent *K. pneumoniae* mediated by mosaic plasmid. (1) Formation of with IncFIB-IncHIB plasmid co-integrate by the fusion of both plasmid replicon (2) Acquisition of virulence plasmid from hvKp strains (3) Formation and maintenance of mosaic plasmid by the integration of virulence associated region into the IncFIB-IncHIB fusion plasmid.

We additionally found that the MDR-hvKp clones carrying mosaic virulence plasmid, possessed *rmpA2* alone without *rmpA*. This observation requires further investigation to determine the mode of acquisition and the stability of *rmpA* and *rmpA2* in this mosaic plasmid. The virulence plasmid found here also carried genes encoding heavy metal resistance, such as tellurium and mercury. Given the community origin of hvKp strains, the co-occurrence of heavy metal resistance is likely to provide an additional survival mechanism in harsh ecological niches (49).

The presence of CRISPR-cas systems in MDR plasmids in *K. pneumoniae* have not been studied extensively. However, recent reports of Type IV CRISPR-Cas system in *K. pneumoniae* mega plasmids/ co-integrate plasmids suggest the role of this system in the competition between plasmids (50, 51). The acquisition of spacers that match with *traL* of conjugative plasmids by *K. pneumoniae* mosaic plasmids suggest the specific targeting of the further invasion of plasmid (51). The attainment of specific plasmid CRISPR spacers targeting different conjugative plasmids appear to be inevitable in *K. pneumoniae* to mitigate the fitness cost associated with carrying multiple AMR plasmids. Notably majority of the plasmids that carried plasmid targeting spacers are co-integrate plasmids carrying IncFIB and IncHI1B replicon (50, 51). This suggest that the plasmid mediated CRISPR spacers not only targets other plasmids but also likely to aid in the formation of co-integrate/ mega plasmids for improved stability and compatibility. The presence of such CRISPR-cas system in our mosaic plasmid further suggest their possible role in the homologous recombination or integration of AMR and virulence determinants in a single plasmid.

Globally, the prevalence of MDR HvKp or CR-HvKp appears to be increasing throughout the past few years. Given the high number of MDR-HvKp and CR-HvKp infections in China, India and Southeast Asia, this region represents the most likely hotspot of MDR virulence overlap and subsequent spread. Similarly, the spontaneous emergence of mosaic plasmids in these regions and its clonal spread in the healthcare setting clearly reflect the burden of these superbugs. If the incidence of the convergent clones with fusion plasmid continues, these pathotypes may be replacing the currently circulating CKPs to become the dominant clones. Since these pathotypes are challenging to treat any further hospital infection outbreak will be fatal.

In India, with its high rates of AMR, it would be a potential healthcare disaster to generate hypervirulent *K. pneumoniae* carrying carbapenemases on a virulence plasmid. The acquisition of ARGs on the chromosome further poses the threat of intrinsic resistance among isolates rendering current empirical a major issue. The convergence of virulence and AMR and the presence of mosaic plasmid are the biggest threats among invasive *K. pneumoniae* infections. It is now apparent that MDR-HvKp isolates are no longer confined to select clones and the containment of such isolates with the mosaic plasmid is very challenging. The presence of AMR and virulence in among diverse *Klebsiella* clones presents a global threat for the rapid spread of these emerging superbugs.

## Author contributions

CS: Conceptualization, Analysis, manuscript writing and revising

KV: Methodology, Bioinformatics, manuscript writing

JJJ: Analysis, Manuscript writing and revising

SB: Manuscript correction and supervision

BJI: Resource

ARN: Methodology, data curation

DPMS: Methodology

BG: Resource

BV: Conceptualization, Manuscript Revision and Supervision

## Ethical approval

The study was approved by Institutional Review Board of Christian Medical College, Vellore, India, with minute number 9,616 (01/09/2015).

---

**Supplementary Figure 1a:**
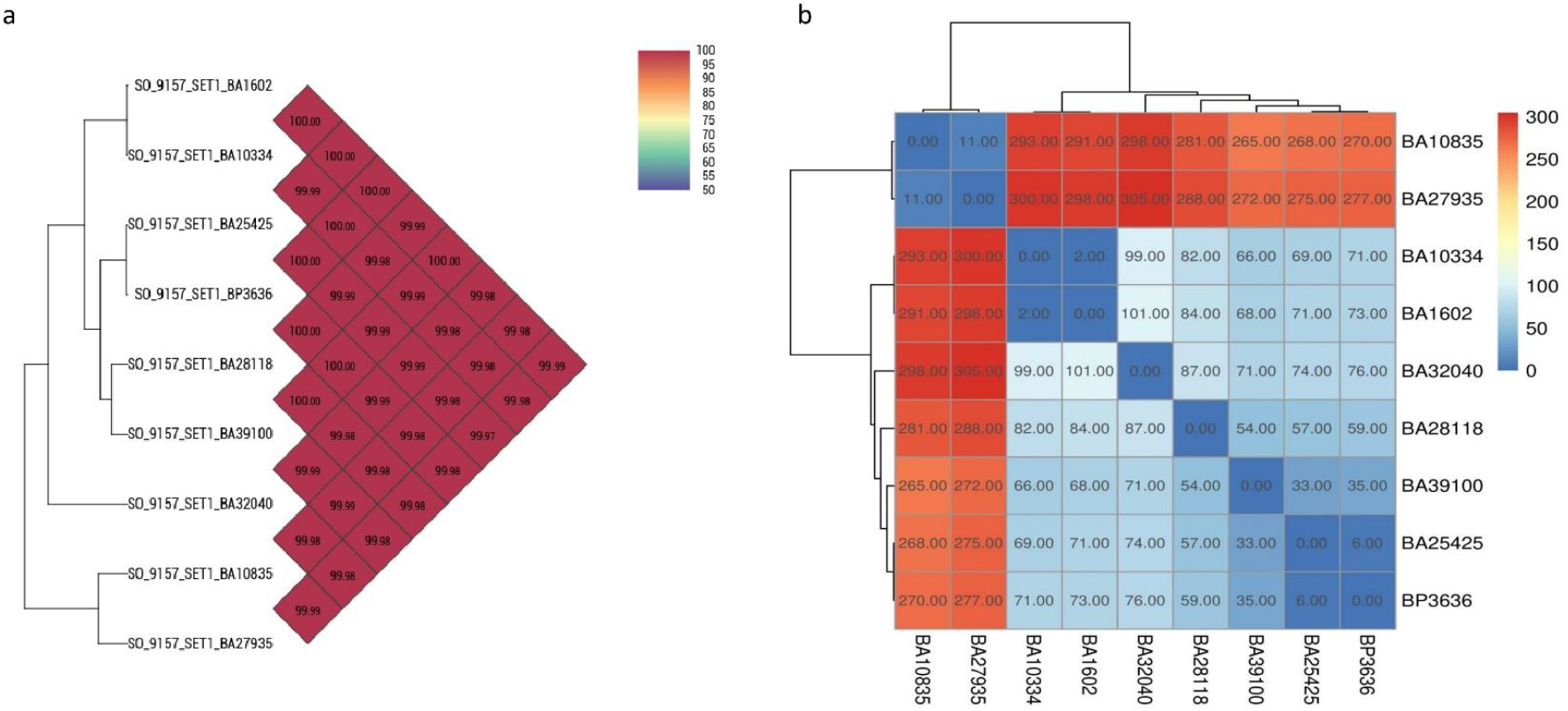
Average nucleotide index (ANI) of nine ST2096 MDR hypervirulent *K. pneumoniae* isolated generated using OrthoANI. Heatmap generated from orthoANI values calculated from the OAT software. The nine study isolates formed two clusters of strains with isolates BA10835 and BA27935 being distinct from the remaining seven. **1b**: Pairwise SNP distance table for all nine sequenced isolates calculated using SNP-dists v 0.6.3 (https://github.com/tseemann/snp-dists). Pairwise SNP difference between the nine study isolates identified two possible clusters of strains with isolates BA10835 and BA27935 being >260 SNP distant from the rest.

